# Profiling tyrosine kinase substrate recognition using bacterial peptide display and deep sequencing

**DOI:** 10.64898/2025.12.28.696772

**Authors:** Minhee Lee, Neel H. Shah

## Abstract

Tyrosine kinases control a wide range of cell signaling pathways that are central to human physiology, and they are dysregulated in a variety of human diseases, most notably cancers. Our understanding of tyrosine kinase biology hinges upon a clear delineation of their protein substrates. Thus, much effort has been invested into defining the substrate specificities of tyrosine kinases, which is partly driven by recognition of the amino acid sequences surrounding the phospho-acceptor tyrosine residues. Numerous methods have been developed to profile tyrosine kinase sequence recognition, and these approaches have collectively demonstrated that different tyrosine kinases have distinct substrate sequence preferences. Here, we describe one such method that combines bacterial peptide display and deep sequencing to study tyrosine kinase substrate preferences. Our approach enables rapid measurement of relative phosphorylation efficiencies for thousands of peptides simultaneously. The genetically-encoded nature of the peptide libraries used with this method allows for facile and cheap construction of libraries tailored to answer a variety questions. Notably, our approach is compatible with genetic code expansion via Amber codon suppression, which allows for the construction and screening of libraries containing non-canonical amino acids. Importantly, results from this assay correlate strongly with quantitative measurements of enzyme kinetics, they corroborate previously reported tyrosine kinase substrate preferences, and they can reveal new insights into tyrosine kinase substrate specificity.

## 1. Introduction

Many cell signaling processes are mediated by the post-translational modification of proteins. Protein tyrosine phosphorylation is one example of a common post-translational modification that is germane to a variety of critical cellular functions in animals, including cell proliferation, differentiation, and apoptosis (Songyang *et al*, 1995). Because of this, aberrant phosphorylation is associated with many human diseases, ranging from cancer to immune diseases to developmental disorders. Tyrosine phosphorylation is controlled by two enzyme families, tyrosine kinases and tyrosine phosphatases. Tyrosine kinase activity is regulated at many levels, including substrate-kinase co-expression and co-localization to controlled protein-protein interactions mediated by non-catalytic domains in both the kinase and substrates (Ubersax & Ferrell, 2007; Miller & Turk, 2018). In addition to these, the catalytic domains of tyrosine kinases are fine-tuned to recognize short linear motifs surrounding the phospho-acceptor residues on their target proteins, which helps mediate selective substrate recognition (Songyang *et al*, 1995) (**Figure 1A,B**). Characterizing the substrate specificity profiles of tyrosine kinases has provided insights into the architectures of cell signaling pathways, the molecular basis for functional specialization across the enzyme family, and mechanisms of pathology of various diseases associated with aberrant tyrosine phosphorylation.

**Figure 1.**
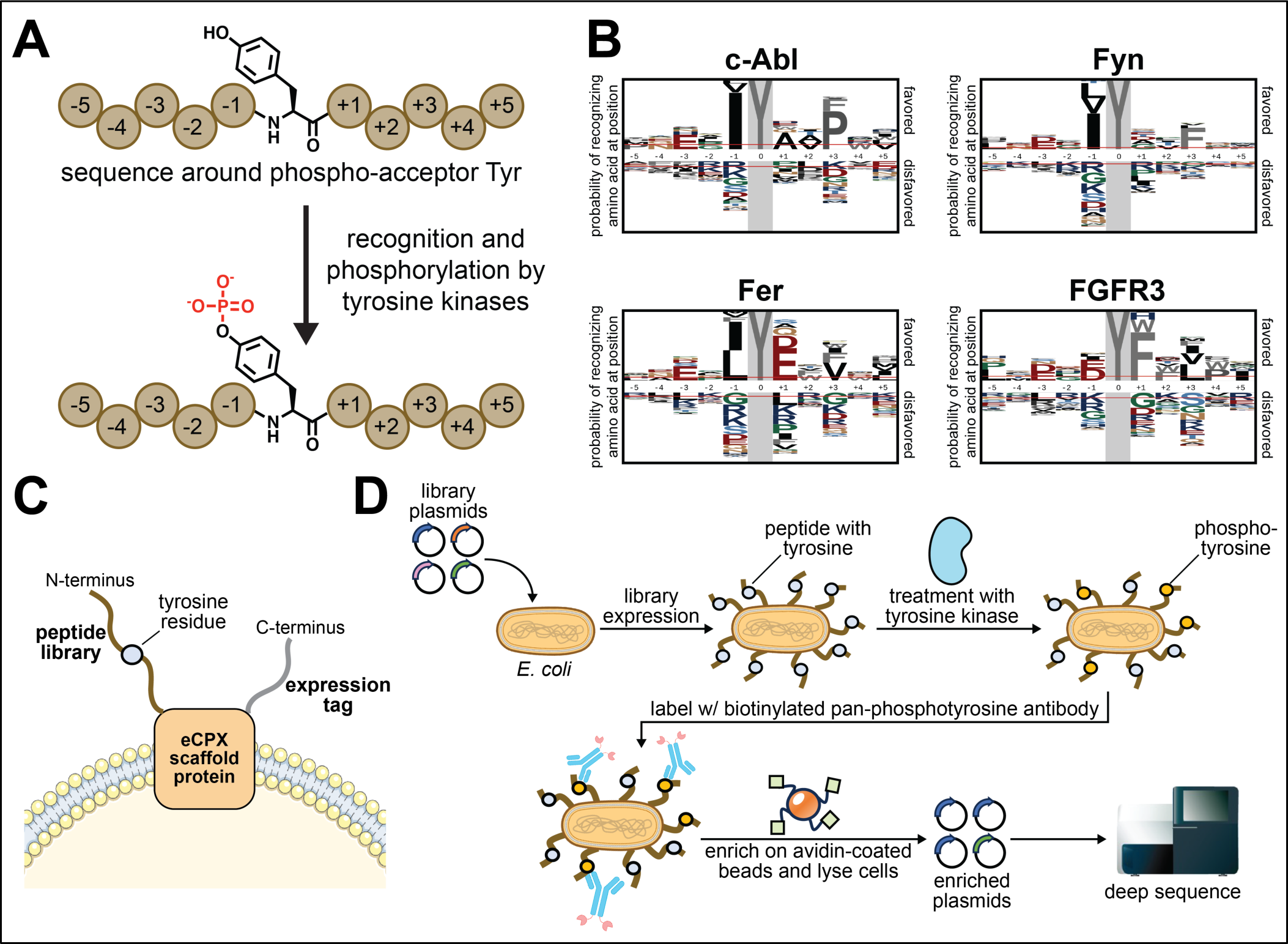
Profiling tyrosine kinase sequence specificity using the bacterial display platform. (**A**) Tyrosine kinases select their substrates, in part, based on recognition of the residues surrounding the phospho-acceptor tyrosine. (**B**) Representative probability logos for various tyrosine kinases, generated using data from the method described here. Logos were generated from selections with a ∼10,000 peptide library, comparing all single-tyrosine peptides with enrichment scores over a cutoff of 3, relative to all single-tyrosine peptides in the library (Li *et al*, 2023). Logos were generated using the pLogo website (https://plogo.uconn.edu/) (O’Shea *et al*, 2013). (**C**) The eCPX scaffold is embedded in the cell membrane and has extracellular N- and C-terminal ends. The peptide libraries are displayed on the N-terminus, and a desired expression tag can be displayed at the C-terminus. (**D**) Bacterial culture is transformed with a genetically-encoded peptide library, allowing for surface display of peptides. The cells are treated with the protein tyrosine kinase of interest, which will preferentially phosphorylate certain displayed peptide sequences. The cultures are treated with a pan-phosphotyrosine antibody conjugated to biotin, allowing for enrichment via avidin-bead pulldown. The cells are lysed and the plasmids are sequenced by deep sequencing for peptide sequence identification and determination of enrichment scores.

Many methods have been developed and reported to characterize tyrosine kinase substrate sequence recognition, each with its own technical merits and drawbacks. The most commonly-used method, pioneered by the Cantley, Turk, and Yaffe labs, involves positional scanning peptide arrays (PSPA), which are arrays with degenerate libraries containing a central tyrosine residue and a fixed residue scanning along the peptide sequence (Songyang *et al*, 1995; Deng *et al*, 2014). In two recent landmark studies, this approach was used to determine the specificity profiles of virtually all human serine/threonine and tyrosine kinases (Johnson *et al*, 2023; Yaron-Barir *et al*, 2024). The resulting position-specific amino acid preferences for each tested kinase can be used to generate high-activity consensus sequences for individual kinases. Furthermore, these data allow for the ranking of preferred substrate sequences across the human proteome. Notably, the comprehensive characterization of a large swath of the enzyme family using one method enhances the power of these predictions, by matching specific phosphorylation sites with the most likely cognate kinase (Johnson *et al*, 2023; Yaron-Barir *et al*, 2024). Overall, PSPAs have proven to be powerful tools for interrogating kinase signaling. One drawback to this approach, however, is ready access to these chemically synthesized libraries. Additionally, because the PSPAs are made up of are degenerate mixtures of peptides, they can only report on position-specific sequence preferences that are independent of sequence context. However, growing evidence suggests that, in some cases, positional preferences can actually depend on the surrounding sequence context (Cantor *et al*, 2018; Li *et al*, 2023).

Other efforts have included strategies to identify the phosphorylation efficiencies of specific sequences, rather than averaged positional residue preferences. One popular example is the one-bead-one-peptide method. This approach has successfully uncovered substrate specificity profiles of various kinases, with the added benefit of reporting discrete substrate sequences (Trinh & Pei, 2016). However, the one-bead-one-peptide method requires the detection, isolation, and individual sequencing of phosphorylated beads, making scale ups to higher throughput technically challenging. Other studies have used defined peptide microarrays, which can include hundreds to thousands of individual sequences immobilized on a solid surface (Warner *et al*, 2008; Amanchy *et al*, 2008; Labots *et al*, 2016). Peptide microarrays can afford higher throughput than one-bead-one-peptide libraries, but the arrays can be expensive. Alternatively, mass spectrometry proteomics-based methods can also reveal kinase sequence preferences. Several groups have expressed kinases of interest in exogenous cell types (e.g. bacteria or yeast), or isolated mammalian cell lysates and treated them with tyrosine kinases, then identified substrates by phospho-proteomics (Xue *et al*, 2012; Sugiyama *et al*, 2019; Finneran *et al*, 2020). Substrates can also be mapped by treatment with highly-selective inhibitors, or using engineered kinases and selective small-molecule modulators, followed by phospho-proteomics (Shah *et al*, 1997; Ferrando *et al*, 2012; Xue & Tao, 2013)12/28/25 6:48:00 PM. These proteomics-based approaches have the benefit that they can reveal substrate preferences in the context of intact proteins, rather than peptides, but results can be complicated by the presence of endogenous kinases or lack of selectivity when using inhibitors.

Molecular display technologies such as bacterial, phage, yeast, and mRNA display have also been used for specificity profiling. In these approaches, genetically-encoded peptide libraries are displayed on a virus/cell surface and phosphorylated, followed by enrichment of phosphorylated cells and sequencing of the peptide-coding DNA. Early versions of these methods faced the bottleneck of relying on Sanger sequencing of individual clones, similar to the challenges of the one-bead-one-peptide approach (Cujec *et al*, 2002; Dente *et al*, 1997). These initial challenges have been since addressed with the advent of deep sequencing (a.k.a. next-generation sequencing) technologies, which enable rapid and quantitative analysis of enrichment across large, complex libraries (Shah *et al*, 2016, 2018; Cantor *et al*, 2018; Taft *et al*, 2019; Li *et al*, 2023). A key benefit to display methods is that they utilize genetically-encoded peptide libraries, which can be made rapidly and cheaply using standard molecular cloning techniques. This allows for the facile construction of bespoke libraries to answer specific mechanistic questions. For example, in our past work using bacterial display and deep sequencing, we have implemented scanning mutagenesis libraries to study the details of molecular recognition for individual sequences (Shah *et al*, 2016; Cantor *et al*, 2018), proteome-derived libraries to profile thousands of physiologically-relevant phosphosites (Shah *et al*, 2018; Cantor *et al*, 2018; Li *et al*, 2023), and larger degenerate libraries that can be used to design consensus sequences and develop predictive models for kinase-substrate recognition (Rube *et al*, 2022; Li *et al*, 2023; Gagoski *et al*, 2025).

Here, we describe a detailed protocol for the use of bacterial peptide display and deep sequencing to profile virtually any protein tyrosine kinase of interest. This method uses an engineered circularly-permuted version of the *E. coli* outer membrane protein OmpX (eCPX), which allows for bi-terminal display of peptide libraries and epitope tags to measure surface display levels (Rice & Daugherty, 2008; Henriques *et al*, 2013) (**Figure 1C**). Prior iterations of this approach relied on the detection of surface-displayed phospho-peptides using fluorescently-labeled pan-phosphotyrosine antibodies, followed by selection using fluorescence-activated cell sorting (Shah *et al*, 2016, 2018; Cantor *et al*, 2018). The streamlined version of this technology (Li *et al*, 2023), described in detail here, utilizes biotinylated pan-phosphotyrosine antibodies and magnetic avidin-functionalized beads as a convenient and higher-throughput alternative to fluorescence-activated cell sorting (**Figure 1D**). Deep sequencing of the bead-enriched libraries allows for robust quantification of enrichment for thousands of individual sequences, and also enables the identification of context-specific sequence preferences. We also describe how this platform can be combined with genetic code expansion techniques to profile kinases against substrate libraries bearing non-canonical amino acids, such as *N*-ε-acetyl-lysine. Overall, the technology described herein provides a unique and approachable strategy to interrogate tyrosine kinase specificity, with the potential to yield novel insights into kinase-substrate recognition that are not accessible using other methods.

## 2. Materials

### 2.1 Construction of genetically-encoded peptide libraries

**Table 1.** Reagents for constructing genetically-encoded peptide libraries.

| Reagent | Additional Notes |
| --- | --- |
| Oligos and primers | See **Table 5** |
| LB agar plates | with antibiotics (diluted 1000-fold from stock solutions) |
| LB media | 25 g Luria Broth powder per 1 L water |
| Antibiotic stocks (1000X) | kanamycin (50 mg/mL), chloramphenicol (25 mg/mL), ampicillin (100 mg/mL), streptomycin (100 mg/mL), all in water, except chloramphenicol, which is dissolved in ethanol. |
| pBAD-eCPX-Myc vector | Any pBAD-eCPX plasmid, bearing any N-terminal displayed peptide sequence or C-terminal expression tag should do, provided the general structure and SfiI sites shown in **Figure 2D** are preserved. |
| SfiI | Restriction enzyme |
| DpnI | Restriction enzyme |
| PB buffer | 5 M guanidinium chloride, 30% (v/v) isopropanol |
| QG buffer | 5.5 M guanidine thiocyanate, 20 mM Tris, pH 6.6 |
| PE buffer | 10 mM Tris, 80% (v/v) ethanol, pH 7.5 |
| 3 M sodium acetate (aq) |  |
| QuickCIP | From NEB |
| 10X CutSmart buffer | From NEB |
| T4 ligase | From NEB |
| 10X T4 ligase buffer | From NEB |
| Electrocompetent 10-beta cells | From NEB |
| Midi or Maxiprep kit | From Zymopure |

### 2.2 Production of recombinant tyrosine kinase domains by bacterial expression

**Table 2.** Reagents for tyrosine kinase domain expression and purification.

| Reagent | Additional Notes |
| --- | --- |
| Plasmid for target kinase domain |  |
| pCDF-YopH | For co-expression with tyrosine kinase domains in bacteria. |
| Electrocompetent or chemically competent BL21(DE3) cells | From NEB |
| LB media | 25 g Luria-Broth powder per 1 L water |
| TB media | 24 g yeast extract, 20 g tryptone, 4 mL glycerol per 900 mL water |
| 10X phosphate buffer | 0.17 M $\text{KH}_2\text{PO}_4$, 0.72 M $\text{K}_2\text{HPO}_4$ |
| 1 M IPTG in water | For inducing protein expression. Working concentration: 0.5 mM. |
| Ni-NTA columns (HisTrap 5mL) | From Cytiva |
| Q-columns (HiTrap 5mL) | From Cytiva, anion exchange column |
| Lysis buffer | 50 mM Tris, 300 mM NaCl, 10 mM imidazole, 10% glycerol (v/v), pH 8.0, 2 mM β-mercaptoethanol (βME) added fresh |
| Wash buffer | 50 mM Tris, 50 mM NaCl, 10 mM imidazole, 10% glycerol (v/v), pH 8.5, 2 mM βME added fresh |
| Elution buffer | 50 mM Tris, 50 mM NaCl, 500 mM imidazole, 10% glycerol (v/v), pH 8.5, 2 mM βME added fresh |
| Anion exchange buffer A | 50 mM Tris, 50 mM NaCl, 10% glycerol (v/v), 1 mM TCEP, pH 8.5 |
| Anion exchange buffer B | 50 mM Tris, 1 M NaCl, 10% glycerol (v/v), 1 mM TCEP, pH 8.5 |
| Kinase SEC buffer | 10 mM HEPES, 150 mM NaCl, 1 mM TCEP, 10% glycerol (v/v), pH 7.5 |
| 100X protease inhibitor cocktail | 20 mM AEBSF dissolved in water, 2 mM leupeptin dissolved in water, 0.1 mM pepstatin A dissolved in methanol |
| Purified TEV protease | can be purified in-house or purchased commercially |
| SDS-PAGE gel | For protein separation |

### 2.3 Bacterial display, selection, and sequencing

**Table 3.** Reagents for bacterial display assay.

| Reagents | Additional Notes |
| --- | --- |
| Luria Broth (LB) |  |
| Electrocompetent MC1061 cells | From Lucigen |
| Electrocompetent 10-beta cells | From NEB |
| Plasmid library |  |
| Antibiotic stocks (1000X) | kanamycin (50 mg/mL), chloramphenicol (25 mg/mL), ampicillin (100 mg/mL), streptomycin (100 mg/mL), all in water, except chloramphenicol, which is dissolved in ethanol. |
| 10% arabinose solution |  |
| Kinase screen buffer | 50 mM Tris, 150 mM NaCl, 5 mM MgCl<sub>2</sub>, pH 7.5 |
| Isolation buffer | PBS, 0.1% BSA (v/v), 2 mM EDTAS |
| QG buffer | 5.5 M guanidine thiocyanate, 20 mM Tris, pH 6.6 |
| PE buffer | 10 mM Tris, 80% (v/v) ethanol, pH 7.5 |
| EDTA solution | 500 mM EDTA, pH 8.0 |
| Dulbecco's phosphate buffered saline (D-PBS) | From Gibco |
| 10% bovine serum albumin (BSA) (w/v) in water | Stored at 4 °C |
| ATP solution | 50 mM in water, pH 7.0 |
| Kinase of interest | Stored in kinase SEC buffer (see **Table 2**) |
| 4G10 Platinum anti-pTyr biotin conjugate | From Fisher |
| Dynabeads™ FlowComp™ Flexi Kit | From Fisher, only the Dynabeads from the kit are needed for this protocol. |
| De-ionized water | (e.g. from a MilliQ water purifier) |
| Oligos and primers | See **Table 5** |
| Q5 Master Mix (2X) for PCR | From NEB |
| TBE buffer | For agarose gel electrophoresis |
| Ethidium bromide | For DNA visualization in agarose gel electrophoresis |
| 1.5% agarose gel | For PCR product separation |
| Quantifluor assay kit for DNA quantification | From Promega |
| Illumina MiSeq Reagent Kit v3 (150 cycles) |  |
| NextSeq 500 Mid-Output v2 Kit (150 cycles) |  |
| Illumina PhiX Control v3 |  |

**Table 4.** Relevant tools and equipment for bacterial display.

| Tools | Additional Notes |
| --- | --- |
| Heated water bath | For bacterial transformation |
| 37 °C incubator | For plate incubation |
| 37 °C incubator with shaking | For liquid culture growth |
| 37 °C and 100 °C heat blocks |  |
| Microcentrifuge | For 1.5-2 mL microcentrifuge tubes |
| Thermocycler | For PCR |
| Gel imager |  |
| Fluorescence plate reader |  |
| Magnetic tube rack for bead separation | During sorting step |
| Illumina MiSeq or NextSeq | Other deep sequencing platforms can also be used, but compatibilities with Illumina adapter sequences (see **Table 5**) must be verified with non-Illumina platforms. |

**Table 5.**
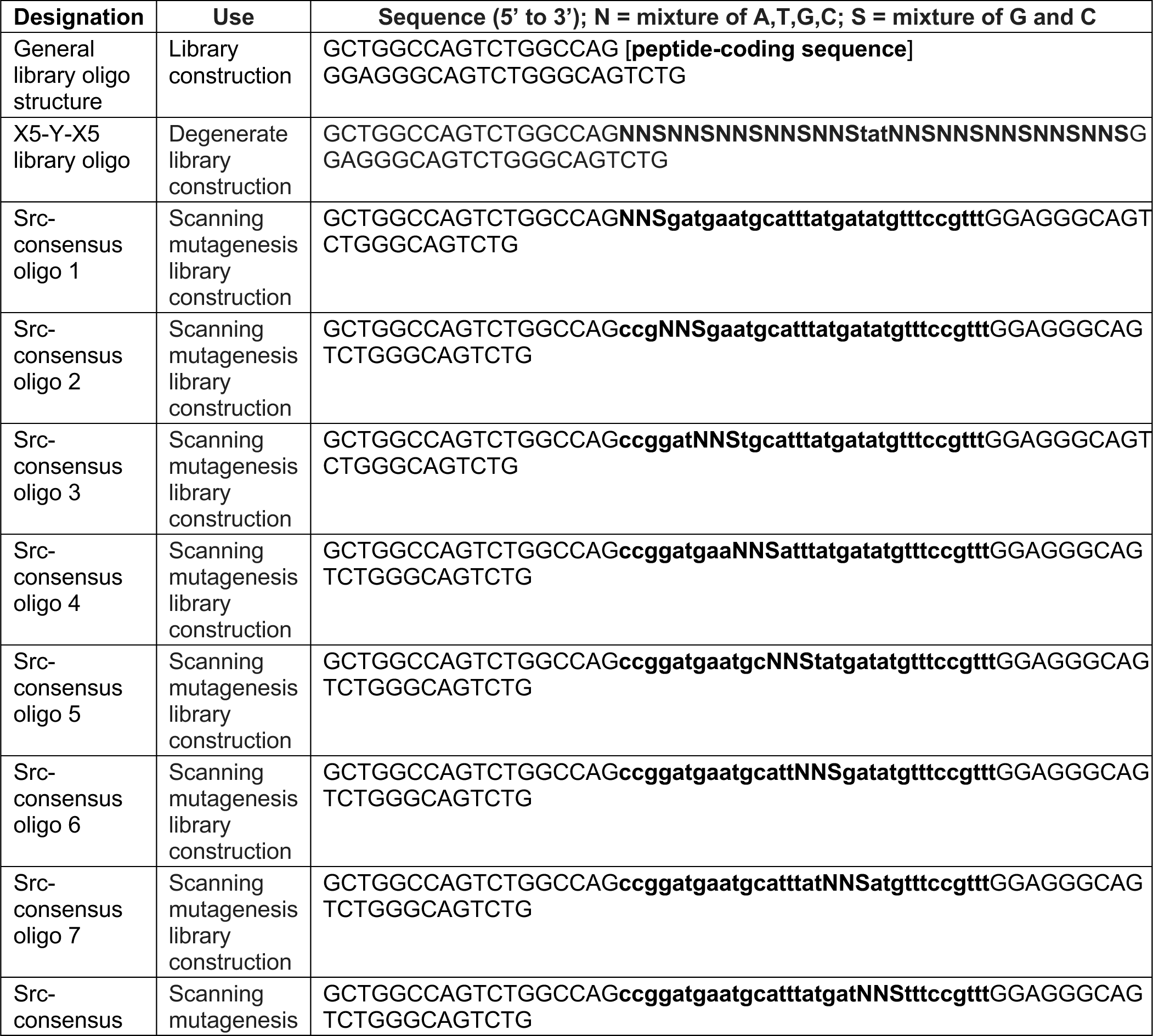
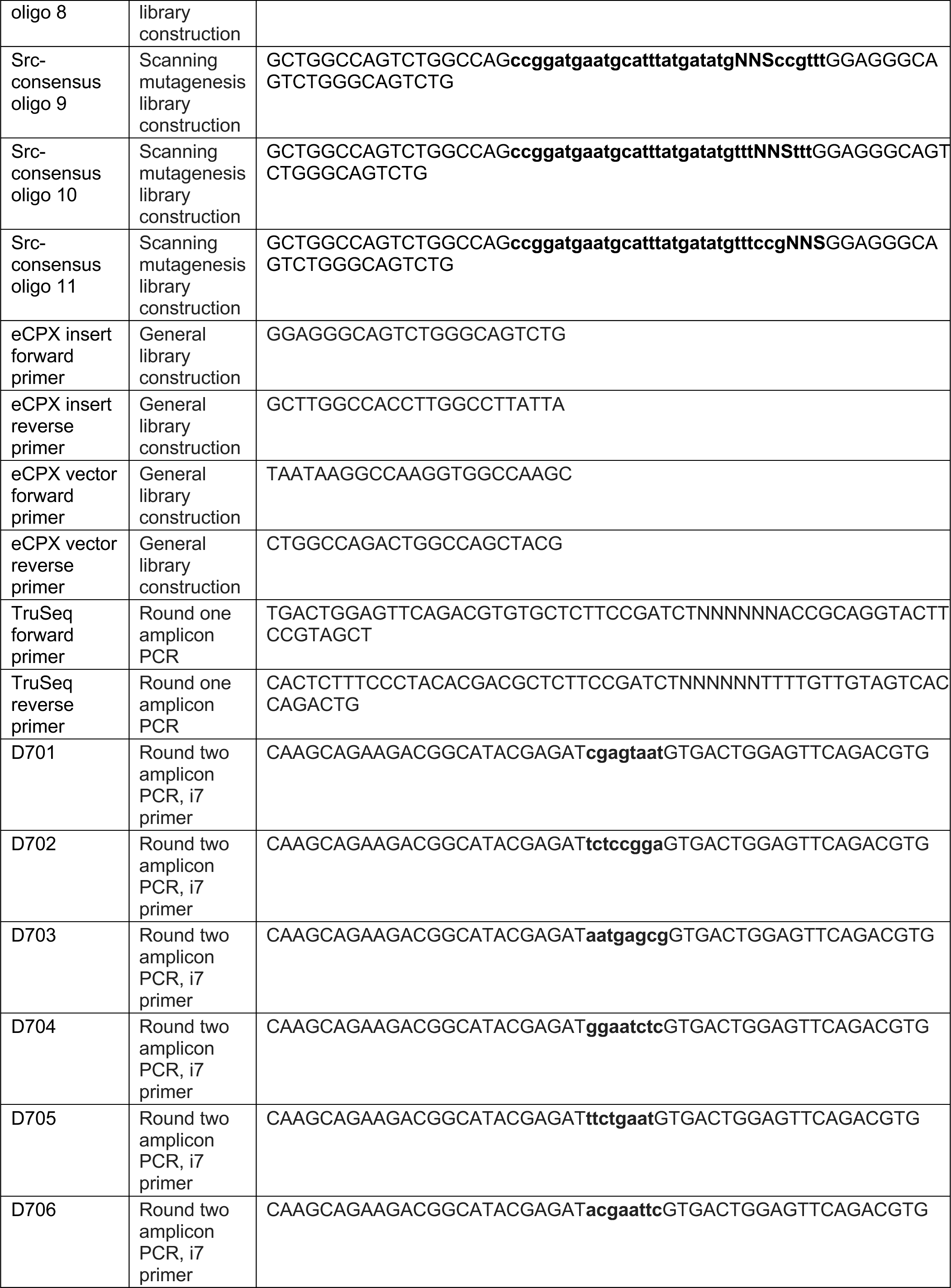
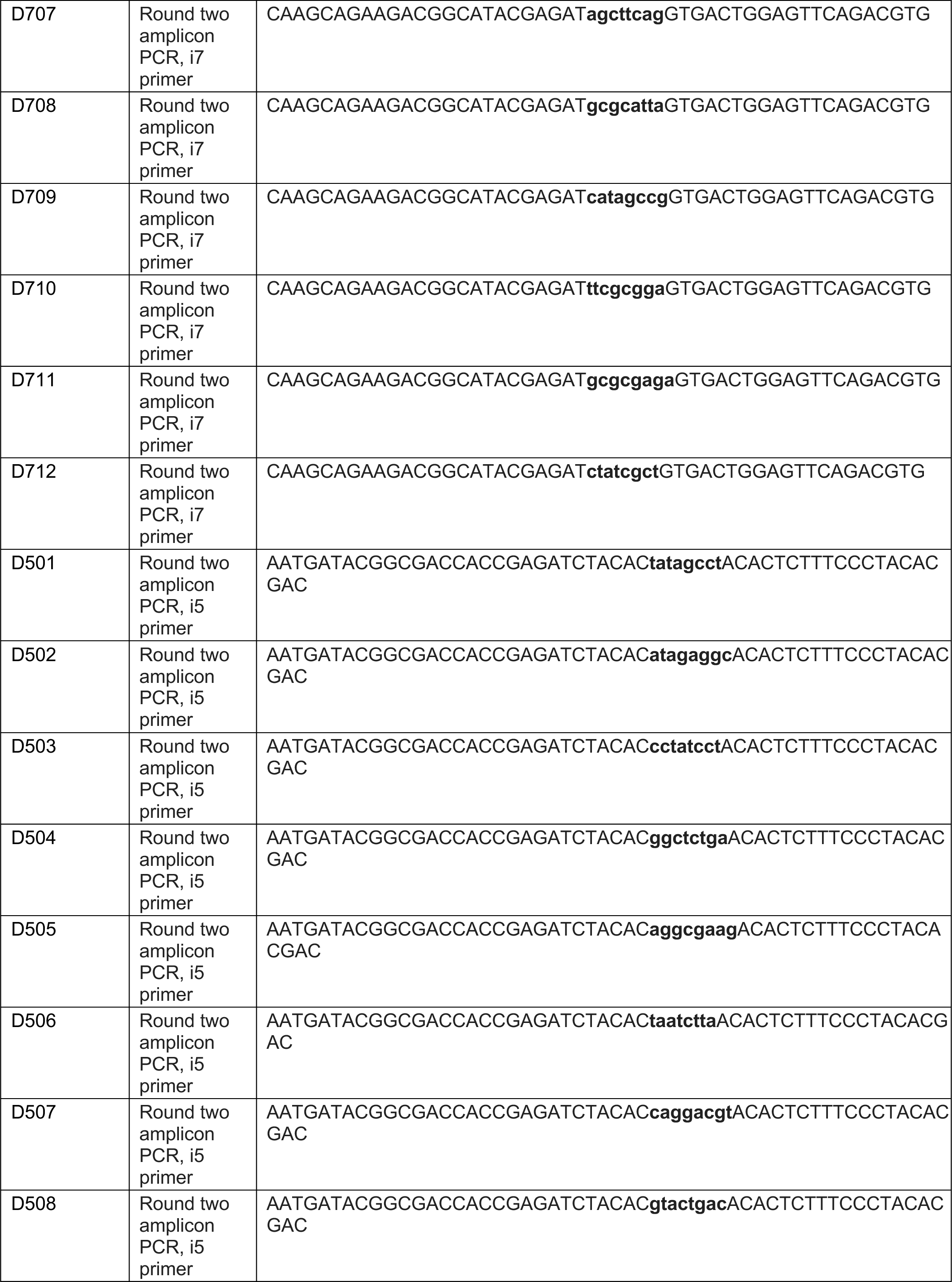
Primer sequences for library construction and downstream sample processing.

## 3. Methods

Carry out all experiments at room temperature, unless otherwise specified.

### 3.1. Construction of genetically-encoded peptide libraries

The method described here is compatible with virtually any genetically-encoded peptide library that is fused to the eCPX scaffold. In our experiments, we have designed and screened three distinct types of libraries: a phospho-site library, a degenerate library, and a scanning mutagenesis library based on an individual substrate sequence (**Figure 2A-C**). We have worked with two phospho-site libraries, designed to encode sequences spanning known human tyrosine phosphorylation sites documented in the PhosphoSitePlus and Uniprot databases (Hornbeck *et al*, 2019; The UniProt Consortium, 2017), along with clinically-observed mutant forms of these sequences, and internal negative controls lacking tyrosine residues. These libraries contain approximately 3,000 and 10,000 sequences, and were generated using synthetic oligonucleotide pools purchased from Twist Biosciences (Shah *et al*, 2018; Li *et al*, 2023). The structure of the oligonucleotides in these pools mirror those used for the degenerate and scanning mutagenesis libraries (**Table 5**). The degenerate library was designed with the degenerate oligonucleotide described in **Table 5**. It encodes sequences with a central phospho-acceptor tyrosine residue with five randomized amino acids flanking both sides of the tyrosine. This library has a theoretical diversity of 20^10^ sequences, which can be expanded further with amber suppression and non-canonical amino acid incorporation. It is noteworthy however, that when accounting for library cloning and bacterial transformation efficiencies, the actual observed diversity of this library is likely closer to 10^6^-10^7^ unique peptide sequences (Li *et al*, 2023; Gagoski *et al*, 2025). This degenerate library enables identification of kinase substrate specificity profiles from an unbiased peptide pool. Scanning mutagenesis libraries are designed based on individual substrate sequences, randomizing each residue to any of the other 19 amino acids. Example oligonucleotides to generate a scanning mutagenesis library based on a Src consensus peptide are given in **Table 5**. These libraries have been used to identify point mutations that either improve or perturb kinase activity, to investigate structure-activity relationships (Shah *et al*, 2016; Cantor *et al*, 2018). Each of these library types can be cloned into the eCPX display vector using the same general strategy described in this section (**Figure 2D,E**).

**Figure 2.**
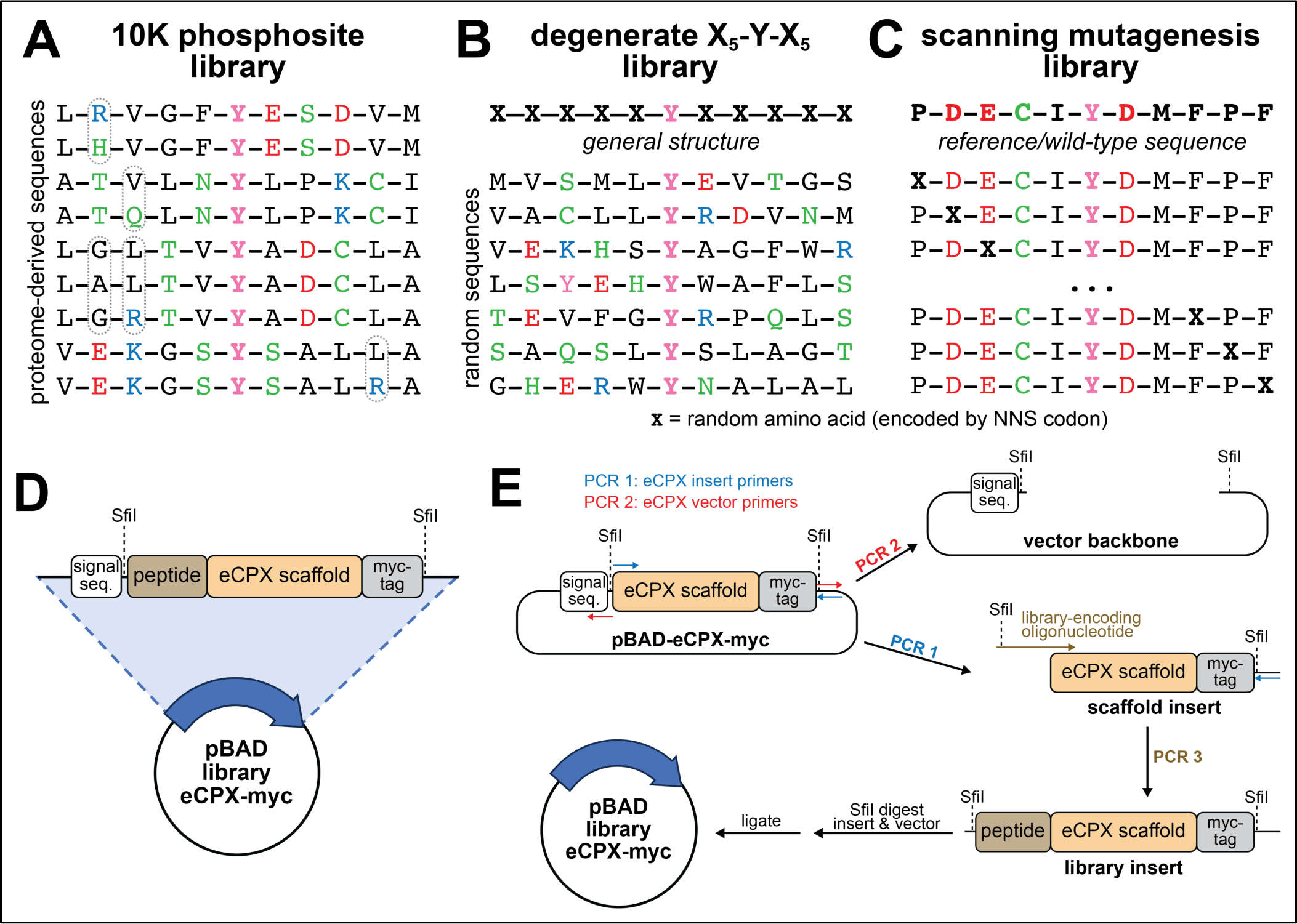
Design and construction of peptide libraries for kinase specificity profiling. (**A**) The 10K phosphosite library, also called the pTyr-Var Library (Li *et al*, 2023), contains a library of defined sequences taken from known phosphosites reported in databases, with clinically-observed mutations and internal negative controls. (**B**) The X_5_-Y-X_5_ degenerate library is a theoretically completely random library in which the “X” positions are encoded by NNS codons and are any one of the 20 canonical amino acids or an amber stop codon. (**C**) The scanning mutagenesis library takes a reference or wild-type peptide sequence and mutates fixed positions to any of the other 19 canonical amino acids (or an amber stop codon) to identify effects of point mutations. (**D**) The completed library plasmids in the pBAD vector have the following elements encoded from the N- to C-terminus: signal sequence, peptide library, eCPX scaffold, Myc tag. (**E**) Cloning approach for generating the genetically-encoded peptide libraries: The “insert region” comprising the eCPX scaffold and Myc tag in the base vector plasmid is amplified out via PCR. In a separate reaction, the vector backbone, including the signal sequence, is amplified out. The insert region undergoes a second PCR to fuse the peptide library to the eCPX scaffold. The completed insert is then cut and pasted into the vector backbone by Sfil digestion and ligation.

#### 3.1.1 General protocol for gel purification of DNA

This section is a general protocol for DNA purification of PCR amplicons, fragments, and/or plasmids from agarose gel following electrophoresis separation, and will be referred to as “***gel purify***”:

1. DNA samples are run on 1-1.5% agarose gels case in TBE buffer and infused with ethidium bromide or an equivalent dye for DNA detection.
2. After imaging the DNA gels under UV light using a gel imager, cut out the desired bands from the agarose gel with a razor. Transfer the cut-out band to a 2 mL microcentrifuge tube.
3. Add 500 μL of QG buffer and incubate at 50 °C until the gel is fully dissolved into solution.
4. Add 10% (v/v) 3 M sodium acetate to the solution and transfer the entire solution to a DNA binding spin column.
5. Centrifuge at max speed (13.3xg) for 1 min. Discard the flow-through and wash with 500 μL QG buffer.
6. Centrifuge at max speed (13.3xg) for 1 min. Discard the flow-through and wash with 750 μL PE buffer.
7. Centrifuge at max speed (13.3xg) for 1 min. Discard the flow-through and repeat the centrifugation step to dry the spin column from ethanol.
8. Transfer the spin column to a 1.5 mL microcentrifuge tube and add 50 μL of pre-warmed deionized water to the spin column. Incubate at RT for 1 min. Centrifuge at max speed (13.3xg) for 1 min.
9. Store the purified DNA sample at -20 °C.

#### 3.1.2 General protocol for PCR cleanup of DNA

This section applies to PCR products following the amplification reactions, and will be referred to as “***PCR cleanup***”:

1. Dilute the total volume of PCR product in 5X volume of PB buffer in a 1.5 mL microcentrifuge tube. Add 3 M sodium acetate in a 1:40 volume ratio (if total volume after adding PB buffer is 480 μL, then 12 μL of 3 M sodium acetate is required).
2. Transfer the PB buffer solutions into separate DNA-binding spin columns, and centrifuge at max speed (13.3xg) for 1 minute. Discard the flow-through and wash the columns with 750 μL PE buffer, centrifuging at max speed for 1 minute. Discard the flow-through and repeat centrifugation to dry the column of all residual ethanol from PE buffer.
3. Transfer the spin column to a 1.5 mL microcentrifuge tube and add 50 μL of pre-warmed deionized water to the spin column. Incubate at RT for 1 min. Centrifuge at max speed (13.3xg) for 1 min.
4. Cleaned DNA samples can be stored at 4 °C for short term storage and at -20 °C for longer time periods.

#### 3.1.3 Construction of libraries

1. The dry primers obtained from vendors can be dissolved in deionized water to a concentration of 100 μM.
2. For the insert and vector PCR reactions, prepare oligonucleotide primer dilutions to 10 μM in deionized water in 1.5 mL microcentrifuge tubes. Combine the forward and reverse primers for each reaction into one tube (eCPX insert primers and eCPX vector primers) mixing in 8 μL of water with 1 μL of each forward and reverse primer.
3. Prepare a PCR reaction for the **insert** with 1X Q5 master mix, 0.5 μM primer mixture, approximately 50 ng of the pBAD-eCPX plasmid vector (template), all diluted up to 50 μL in deionized water.
4. Prepare the PCR reaction for the **vector backbone** in the same way as the insert reaction, but with a total reaction volume of 200 μL. Divide this into 4 equal aliquots of 50 μL to generate 4 vector backbone reactions.
5. Run the PCR according to the following parameters for the insert region (**Figure 2E)**: Step 1: 98 °C, 30 sec Step 2: 98 °C, 10 sec Step 3: 62 °C, 10 sec Step 4: 72 °C, 30 sec Step 5: Return to Step 2, 24 cycles Step 6: 72 °C, 2 min Step 7: 4 °C, hold
6. Run the PCR according to the following parameters for the vector backbone region (**Figure 2E**): Step 1: 98 °C, 30 sec Step 2: 98 °C, 10 sec Step 3: 71 °C, 10 sec Step 4: 72 °C, 165 sec Step 5: Return to Step 2, 26 cycles Step 6: 72 °C, 3 min Step 7: 4 °C, hold
7. ***Gel purify*** the cloned **insert** fragments on 1.5% agarose gel, separating at 140 V (target base pairs for insert = 545 bp), as described.
8. For the vector backbone fragments, add 1 μL of DpnI restriction enzyme to each PCR reaction, vortex, and spin down in a centrifuge. In a thermocycler, incubate the reactions at 37 °C for 1 hr, followed by a 80 °C heating step for 20 min, with a final hold at 4 °C until the next step.
9. For the 2nd round of PCR for the insert, the peptide library sequences will be fused to the insert region (**Figure 2E**). Prepare a primer mix as described in step 2, containing the eCPX insert reverse primer with the appropriate library primer (or oligo pool) at 10 μM.
10. Prepare the PCR reaction for this **second insert reaction** with 4 μL of the primer mix, 1 μL of the purified eCPX insert (from the first PCR, roughly 50 ng of DNA), 40 μL of 2X Q5 master mix, all diluted up to 80 μL in deionized water. Divide the total reaction volume into 10 μL aliquots (there should be 8x10 μL reactions). Note that splitting the PCR reaction at this step could limit PCR bias in the library. In cases where individual sequences in the pool get preferentially amplified during early cycles, individual sub-reactions may not contain the same biases.
11. Run the PCR according to the following parameters for the insert region: Step 1: 98 °C, 30 sec Step 2: 98 °C, 10 sec Step 3: 68 °C, 10 sec Step 4: 72 °C, 30 sec Step 5: Return to Step 2, 24 cycles Step 6: 72 °C, 2 min Step 7: 4 °C, hold
12. Pool the eCPX-insert reaction products (80 μL total) into a 1.5 mL microcentrifuge tube, and do a ***PCR cleanup***. Pool the backbone reaction products (200 μL total) into a separate 1.5 mL microcentrifuge tube, and do a ***PCR cleanup***.
13. Prepare Sfil digestion reactions for the cleaned eCPX-insert and backbone vector PCR products: **Insert**: 25 μL of 10X CutSmart buffer, 50 μL of purified eCPX-insert, 12 μL Sfil, all diluted up to 250 μL total with deionized water. Divide the total volume into 5x50 μL aliquots. **Vector backbone:** 20 μL of 10X CutSmart buffer, 100 μL of purified vector backbone, 16 μL Sfil, all diluted up to 200 μL total with deionized water. Divide the total volume into 4x50 μL aliquots.
14. Incubate the Sfil reactions at 50 °C overnight (16-18 hours). These products can be stored at 4 °C for longer term storage.
15. The vector backbone must be further treated for 5’-end dephosphorylation required for efficient DNA ligation. After the Sfil reaction incubation, add 2 μL of QuickCIP to each PCR reaction and incubate at 37 °C for 30 min.
16. Following the QuickCIP reaction for the vector backbone products and the Sfil reaction for the eCPX-insert products, ***gel purify*** the products on 1% agarose gel. Elute each product in 50 μL of deionized water, and pool together like-samples as necessary.
17. Measure the concentration of the purified products by Nanodrop or equivalent method.
18. The eCPX-insert and vector backbone products will now be ligated to reconstitute the plasmid that genetically encodes the eCPX scaffold for bacterial peptide display, as well as the peptide libraries of interest (**Figure 2E**). For each desired library, add 2.4 μg of vector backbone product, 1.28 μg of eCPX-insert product, 40 μL of T4 ligase, 80 μL of 10X T4 ligase buffer, all diluted up to 800 μL in deionized water.
19. Divide the total volume into 16x50 μL aliquots into PCR tubes. Incubate the reactions at 16 °C overnight (16-18 hours), followed by a 80 °C denaturation step for 20 min, finished by a 4 °C holding step for short-term storage.
20. Pool the ligation products into a 1.5 mL microcentrifuge tube, and do a ***PCR cleanup***. However, during the elution step, elute in 30-40 μL of deionized water to obtain a higher DNA concentration.
21. Measure the concentration of the cleaned ligation products by Nanodrop or equivalent method.
22. Thaw a 100 μL aliquot of electrocompetent 10-beta cells on ice and add 1 μg of ligated product. Electroporate the cells at 1.80 kV in a 1 mm pathlength electroporation cuvette, and recover the cells in 950 μL of pre-warmed 10-beta cell media. Recover the cells at 37 °C, rotating at 215 rpm for 1 hour.
23. Following the recovery incubation, prepare 10-fold serial dilutions of the recovery culture in 1.5 mL microcentrifuge tubes by removing 10 μL from the recovery culture and diluting in 90 μL media, then further diluting 10 μL of this into 90 μL media, then doing this one more time, for 10-, 100-, and 1000-fold dilutions.
24. Plate 80 μL of each serial dilution onto LB agar plates containing 25 μg/mL chloramphenicol. Incubate the plates overnight (16-18 hours) at 37 °C for colony growth.
25. Add the rest of the recovery culture to 250 mL of LB media containing 25 μg/mL chloramphenicol and incubate overnight (16-18 hours) at 37 °C, rotating at 215 rpm.
26. Count the number of cells that are countable for the serial dilution plates to estimate the number of transformants on that plate, then multiply by the dilution factor to approximate the number of transformants in the whole transformation mixture. The goal is to have >100 transformants per variant for smaller phosphosite and scanning mutagenesis libraries, and at least 10^6^-10^7^ transformants for degenerate libraries (although 10^8^ is desirable).
27. Divide the 250 mL of overnight growth culture into 5x50 mL conical tubes, and centrifuge at 4000g for 15 min to pellet the cells. Discard the supernatant. The pellets can be stored at -80 °C for long-term storage.
28. Extract the cloned DNA plasmid library by Midiprep (Zymopure), eluting in 200 μL of deionized water.
29. Determine the library DNA concentration by Nanodrop or equivalent method and store the DNA at -20 °C or -80 °C for long-term storage.

### 3.2. Production of recombinant tyrosine kinase domains by bacterial expression

This kinase profiling method relies on the use of purified kinases. For most of our experiments, we have used isolated tyrosine kinase domains, as they omit contributions from non-catalytic domains (e.g. SH2 domains that bind to specific tyrosine-phosphorylated sequences) that could confound interpretation of the results. We often express and purify the kinase domains from *E. coli* (Li *et al*, 2023), but other sources of active enzymes will also work, including commercially purchased kinase constructs, or kinases expressed from other cell types (e.g. insect or mammalian cells) (Shah *et al*, 2016). In some cases, isolated kinase domain constructs are not sufficiently active, and further construct engineering may be necessary. For example, EGFR kinase requires the formation of an asymmetric dimer between two EGFR kinase domains for sufficient activity. This was previously achieved using rapamycin-inducible EGFR kinase domain constructs fused to FKBP and FRB, yielding a construct that was compatible with phosphorylation of the bacterial display platform (Cantor *et al*, 2018). Alternatively, other kinases may require pre-phosphorylation to achieve sufficient activity levels. For example, FGFR1 kinase requires a pre-incubation step with ATP followed by desalting to activate the kinase domain for use in the display platform (Li *et al*, 2023). Here, we describe our standard protocol for the purification of kinase domains from bacteria.

1. Take an aliquot of BL21(DE3) electrocompetent cells and add 1 μL each of pCDF-YopH-StrepR and the plasmid with the target His_6_-tagged tyrosine kinase domain (IPTG-inducible, typically with ampicillin or kanamycin resistance).
2. Electroporate at 1.80 kV in a 1 mm pathlength electroporation cuvette, and recover in 1 mL of pre-warmed LB media. Transfer the culture to a 1.5 mL microcentrifuge tube and incubate at 37 °C for 1 hour, shaking at 215 rpm.
3. Plate 100 μL of the cells on LB agar plates, with corresponding antibiotic (streptomycin for the YopH plasmid and the appropriate antibiotic for tyrosine kinase plasmid).
4. Incubate the plates at 37 °C for overnight (16-18 hours).
5. Around late afternoon the next day, pick a colony and inoculate 100 mL of LB media with corresponding antibiotics in a 250 mL Erlenmeyer flask to generate the starter culture. Incubate at 37 °C, shaking at 215 rpm overnight (16-18 hours).
6. The following day, dilute the starter culture to OD_600_ 0.1 in 1 L of TB media treated with appropriate antibiotics. Incubate at 37 °C, shaking at 215 rpm until OD_600_ reaches about 0.5-0.6.
7. Once the OD_600_ reaches 0.5-0.6 (mid-log phase), add 0.5 mM IPTG to induce protein expression and incubate at 18 °C, shaking at 215 rpm overnight (16-18 hours).
8. The next day, centrifuge the cultures at 4000g for 30 min at 4 °C. Carefully decant the media supernatant.
9. Add 7 μL of βME to 50 mL of lysis buffer (final concentration of 2 mM βME) and resuspend the cell pellets in 15 mL of this solution. Add 150 μL of 100X protease inhibitor cocktail (**Table 2**).
10. Once fully resuspended, transfer the cells to a 100 mL beaker and sonicate on ice using a probe sonicator to lyse (other lysis methods can also be used here).
11. Transfer the lysate to a 50 mL tube and centrifuge at 14,000g for 45 min to pellet debris.
12. Meanwhile, equilibrate the Ni-NTA and Q-columns by running deionized water for 10 min, followed by lysis buffer and anion exchange buffer A, respectively, for another 10 min, flowing at 2-3 mL/min minutes using a peristaltic pump.
13. Place the Q-column on ice after the equilibration step.
14. Filter the centrifuged lysate through a 1.1 μm glass fiber filter and load onto the Ni-NTA column using the peristaltic pump. Collect the flow-through in a conical tube. Also save a small gel sample (∼15 μL) of the filtered lysate and flow-through for subsequent analysis.
15. Wash the column with 50 mL of lysis buffer with 7 μL βME. Collect the flow-through in a conical tube and store on ice.
16. Wash the column with 50 mL of wash buffer with 7 μL βME. Collect the flow-through in a conical tube and store on ice.
17. Stack the Q-column after the Ni-NTA column (flow-through from nickel column should be flowing directly into the Q-column) and wash with 50% wash/elution buffer with 7 μL βME. Collect the flow-through in a conical tube and store on ice.
18. Remove the Ni-NTA column and attach the Q-column to the peristaltic pump. Wash the column with 50 mL of anion exchange buffer A. Collect the flow-through in a conical tube and store on ice.
19. Attach the Q-column to an FPLC pump and run over a linear gradient from 100% anion exchange buffer A to 100% anion exchange buffer B at 1 mL/min over 10 column volumes. Collect 5 mL fractions across the whole gradient and store on ice or at 4 °C.
20. Take 15 μL from all collected flow-through samples and fractions for SDS-PAGE analysis.
21. Run all samples on an SDS-PAGE gel and Coomassie stain the gel for visualization.
22. Based on the gel, consolidate the desired fractions eluted from the FPLC and set aside 15 μL for gel analysis. Add TEV protease for TEV cleavage and incubate at RT for 2 hours and store at 4 °C overnight. The exact quantity of TEV, temperature, and duration of the reaction may need to be optimized, depending on the specific construct.
23. Reserve 15 μL of the post-TEV solution for gel analysis.
24. Load the solution onto an Ni-NTA column pre-equilibrated in lysis buffer and collect the flow-through in a conical tube, stored on ice.
25. Wash the column with 3x 10 mL lysis buffer (with 3 μL βME). Collect the flow-through in a conical tube and store on ice.
26. Add 20 mL of 50% wash/elution buffer (with 5 μL βME) to the column to elute any bound proteins. Collect the flow-through in a conical tube and store on ice.
27. Take 15 μL from all collected flow-through and elution fractions and analyze by SDS-PAGE.
28. Based on the gel, consolidate the desired fractions and concentrate to 1-3 mL using a centrifugal filter, keeping the protein cold the whole time.
29. Load the sample onto a size exclusion chromatography column, running with kinase SEC buffer. Collect appropriate-sized fractions (0.5 mL to 1 mL) throughout the run.
30. Take 15 μL from all SEC fractions and analyze by SDS-PAGE with Coomassie staining.
31. Based on the gel, consolidate the desired fractions and concentrate to σ:100 μM. Estimate protein concentration using a colorimetric assay (e.g. Bradford) or via absorbance at 280 nm.
32. Prepare aliquots (σ:50 μL), flash-freeze in liquid nitrogen, and store at -80 °C.

### 3.3 Expression of bacterial peptide display libraries

The bacterial display method relies on the expression of the eCPX scaffold with attached peptide library and c-Myc (or other) expression tag on the cell surface. *E. coli* strain MC1061 cells work well for this purpose and the workflow presented will be largely analogous to the protein expression procedure described above. **Section 3.5** will discuss key changes to include non-canonical amino acids during this step. Here, we describe our standard protocol for the expression of eCPX scaffold and peptide libraries in MC1061 bacterial cells:

1. Thaw a 100 μL aliquot of electrocompetent MC1061 cells on ice. Add 1 μL (∼100 ng) of the desired constructed plasmid library to the culture aliquot and mix by very gently pipetting up and down, or by flicking the tube.
2. Electroporate at 1.80 kV in a 1 mm pathlength electroporation cuvette, and recover the cells in 1 mL of pre-warmed LB media. Transfer the culture to a 1.5 mL microcentrifuge tube and incubate at 37 °C for 1 hour, rotating at 215 rpm.
3. Dilute the entirety of the recovery culture in 50 mL of LB, containing 25 μg/mL chloramphenicol. Incubate at 37 °C, rotating at 215 rpm for overnight.
4. The next day, dilute 150 μL of the overnight culture into 5 mL of LB containing 25 μg/mL chloramphenicol, and incubate at 37 °C, rotating at 215 rpm until OD_600_ reaches 0.5-0.6.
5. Once the target OD_600_ is reached, induce expression with 0.4% arabinose (v/v) and incubate at 25 °C for 4 hours, rotating at 215 rpm.
6. After induction, centrifuge the cultures at 4000g for 10 min to pellet. Decant the media supernatant and gently resuspend the cells in 5 mL of cold D-PBS. The resuspended cells can immediately be used in the next step or stored temporarily (see **note 1**).

### 3.4 Running the bacterial peptide display selection assay

Once the bacterial cells have been prepared with expressed peptide library, the cells can be treated with the target tyrosine kinase of interest. The kinase will preferentially phosphorylate the displayed peptide bearing optimal sequence features. The cells displaying phosphorylated peptides can then be labeled with a biotinylated pan-phosphotyrosine antibody and enriched by magnetic avidin-coated bead. The protocol described below discusses our standard procedure for sorting peptide libraries following library expression and phosphorylation:

1. Prepare 150 μL aliquots from the resuspended cultures in 1.5 mL microcentrifuge tubes. Centrifuge at 4000g for 10 min to pellet, and carefully remove the PBS supernatant.
2. Meanwhile, prepare a working kinase screen buffer by adding 2 mM sodium orthovanadate and 1 mM TCEP to a desired volume of the kinase screen buffer, based on the number of parallel phosphorylation reactions to be carried out (see next step).
3. After removing the PBS supernatant, resuspend the cells in 100 μL of the working kinase screen buffer (with orthovanadate and TCEP added).
4. Add a desired concentration of purified tyrosine kinase domain to the samples. Alternatively, the kinase can be pre-diluted in the working kinase buffer first, then this solution can be used to resuspend the cells. Kinase concentrations used in past experiments are given in **Table 6**.
5. Add a final concentration of 1 mM ATP to the samples to start the kinase phosphorylation reactions and mix gently. Incubate at 37 °C for a desired amount of time. The reaction conditions used for our experiments are listed in **Table 6**. *As stated in step 4, the kinase concentration and incubation times will have to be optimized for the desired level of phosphorylation beforehand for other unlisted kinases (see **note 2**)*.
6. Quench the reactions upon completion with a final concentration of 25 mM EDTA, added from a 500 mM EDTA, pH 8.0, stock solution. Centrifuge the samples at 4000g for 10 min to pellet.
7. Remove the supernatant and wash the cells in 200 μL of cold D-PBS/0.2% BSA (v/v) and repeat the centrifugation to remove the supernatant.
8. In the meantime, during the wash step, prepare a 1:1000 dilution of anti-phosphotyrosine 4G10 Platinum biotinylated antibody in PBS/0.2% BSA. Prepare enough volume for 100 μL per sample.
9. Resuspend the washed pellets in 95 μL of the antibody solution and incubate on ice for 1 hour.
10. After incubation, wash the cells as described in step 7 with 200 μL of cold PBS/0.1% BSA. **In this step before centrifugation, you can discard half of the volume (100 μL) to reduce your working sample volume, if desired (see note 3). If this is the case, reduce all volumes listed below until step 16 by half unless otherwise specified.**
11. Resuspend the washed cells in 1 mL of isolation buffer (PBS, 0.1% BSA, 2 mM EDTA).
12. To prepare the magnetic Dynabeads for use, measure out enough volume for 75 μL/sample into a microcentrifuge tube and place on a magnetic rack. Remove the stock buffer and wash the beads in 1 mL of isolation buffer. Repeat the separation step to remove the buffer, and resuspend the beads in the *same initial volume* with isolation buffer.
13. Once the beads have been washed, add 75 μL beads to each sample and incubate at 4 °C for 20 min, rotating gently to mix.
14. Place the samples on magnetic racks to isolate the beads and bound cells on the side of the tube. Remove the supernatant and wash with 1 mL of isolation buffer for each sample. Incubate at 4 °C for 15 min, rotating gently to mix.
15. Repeat the magnetic bead separation step and remove the isolation buffer. Add 50 μL of fresh deionized water to each sample to resuspend the beads and bound cells (elute in 50 μL deionized water even if half bead volumes were used). Boil these samples for 10 min on a heat blot (see **note 4**). The samples will now be lysates.
16. Centrifuge the samples at max velocity in a microcentrifuge for 10 min at 4 °C to pellet the beads. The lysates can be stored at -20 °C.
17. To prepare an unsorted reference lysate, take 150 μL of the resuspended cultures from each transformant. Centrifuge at 4000g for 10 min to pellet the cells and remove the supernatant.
18. Resuspend these cells in 300 μL of deionized water and boil for 10 min.
19. Centrifuge the samples at max velocity for 10 min at 4 °C to pellet the cell debris. These unsorted lysates can be stored at -20 °C, alongside the sorted samples.

**Table 6.** Reaction conditions for kinase phosphorylation of displayed peptide libraries.

| Kinase domain | Kinase concentration (µM) | Incubation time at 37 °C (min) |
| --- | --- | --- |
| c-Src | 0.5 | 3 |
| c-Abl | 1.5 | 3 |
| Fer | 0.4 | 3 |
| EPHB1 | 1.5 | 3 |
| EPHB2 | 1.25 | 3 |
| JAK2 | 0.1 | 3 |
| AncSZ | 0.5 | 3 |
| FGFR1* | 0.45 | 3 |
| FGFR3* | 0.5 | 3 |
| MERTK* | 0.7 | 3 |

\* The marked kinases were pre-activated with 25 µM kinase domain and 5 mM ATP for 0.5-2 hours at 25 °C.

### 3.5 Combining bacterial peptide display with amber suppression

As described above in **section 3.3**, this bacterial display platform is also compatible with amber suppression for the incorporation of non-canonical amino acids. To incorporate non-canonical amino acids such as *N*-ε-acetyl-lysine, we use the pUltra vector encoding for the corresponding aminoacyl synthetase and tRNA. However, because the MC1061 cells and pUltra vector have overlapping streptomycin resistance, the pUltra plasmids were modified to have ampicillin resistance instead, which is orthogonal to the resistances of the pBAD-eCPX vector (chloramphenicol) and MC1061 cell line (streptomycin) (**Figure 3**). This can be achieved for any pUltra vector by simply swapping the streptomycin resistance gene coding sequence with an ampicillin resistance gene coding sequence from any other plasmid, using Gibson assembly or any other restriction enzyme-free cloning method. Finally, we note that libraries containing degenerate ‘NNS’ codons (e.g. the X5-Y-X5 library and scanning mutagenesis libraries) contain sequences bearing an amber codon (TAG), and are thus compatible with the approach described below, without further library engineering. In this section, we describe the key changes in the protocol listed in **section 3.3** to accommodate non-canonical amino acid incorporation once the orthogonal plasmid is acquired:

**Figure 3.**
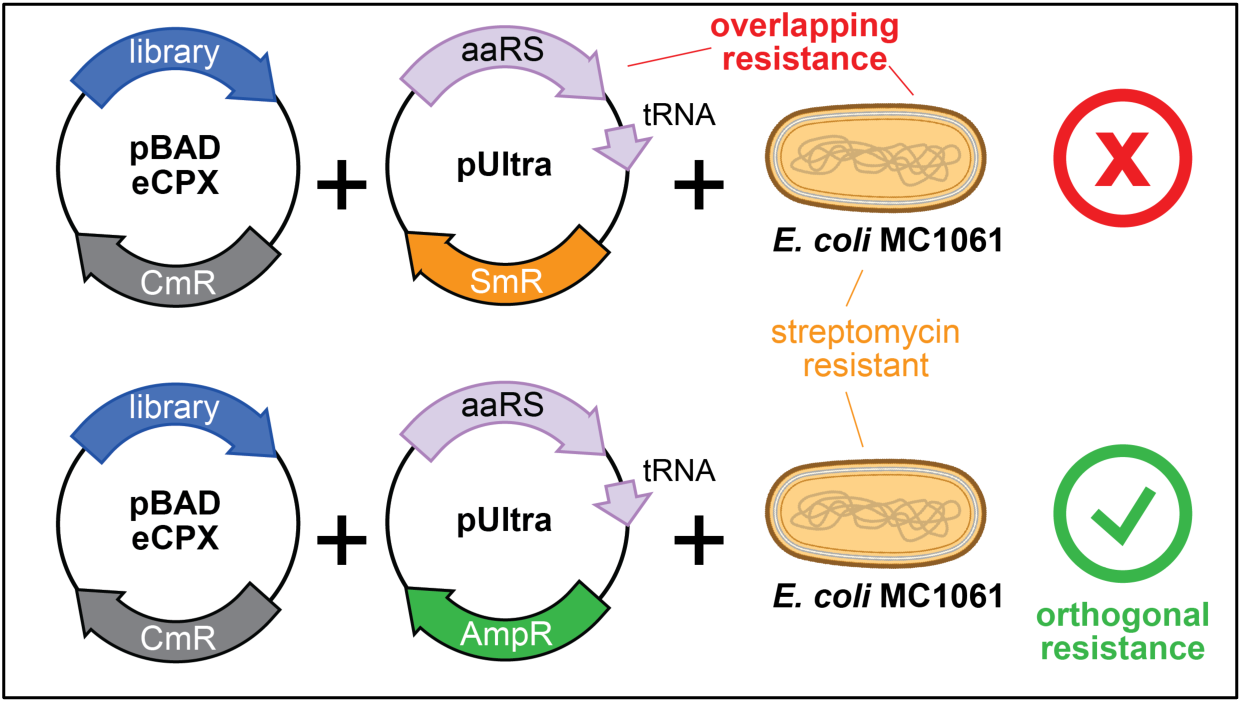
Orthogonal antibiotic resistance for non-canonical amino acid incorporation via amber suppression. The peptide library is encoded on the pBAD plasmid, which has a chloramphenicol resistance marker (CmR). The display experiments are typically done in *E. coli* MC1061 cells, which have intrinsic streptomycin resistance (SmR). A commonly-used plasmid encoding aminoacyl tRNA synthetases (aaRS) and suppressor tRNA molecules for non-canonical amino acid incorporation, the pUltra plasmid, also has streptomycin resistance, making it incompatible with MC1061 cells. To address this, we replaced the streptomycin resistance gene with ampicillin resistance, ensuring that all components have orthogonal resistances.

#### Repeat the same steps as listed above in section 3.3 with the following adjustments. The numbering is intended to match the corresponding steps in section 3.3

1 Add 1 μL of the plasmid with the corresponding synthetase and tRNA of the non-canonical amino acid of interest, to the cell aliquot alongside the library plasmids.
4 In addition to 25 μg/mL chloramphenicol in the growth media, also add 100 μg/mL ampicillin.
5 Alongside 0.4% arabinose during the induction step, add the desired concentration of non-canonical amino acid, and other inducing agent required for expression of aminoacyl synthetase and tRNA (eg: IPTG for pUltra vectors).

We have used pUltra-AcKRS plasmid for our synthetase and tRNA, and induced expression with 0.5 mM IPTG and 10 mM acetyl lysine (AcK).

*The amber suppression parameters will have to be optimized beforehand for best incorporation and expression efficiencies for other desired non-canonical amino acids*.

### 3.5 Preparing sequencing samples from the selected cells

After preparing the unsorted and sorted lysates, the library segments must be amplified and purified for deep sequencing analysis. The oligos listed in **Table 5** will be used to attach TruSeq adapter sequences and the barcode i7/i5 sequences to the library-encoding regions by PCR. High-fidelity methods must be used to determine DNA concentrations following post-PCR purification for accurate loading and to avoid over- or under-clustering during deep sequencing. The protocol listed below describes our standard procedure for processing and quantifying DNA samples after sorting.

#### 3.5.1 Amplifying the DNA from the lysates after library phosphorylation and selection

1. Prepare a primer mix with the TruSeq primers, diluting each down to 5 μM in water.
2. Prepare a PCR reaction for each sample with the following volumes: 20 μL lysate (be sure to include a sample with the unsorted lysate) 5 μL TruSeq primer mix 25 μL 2X Q5 master mix
3. Run the PCR according to the following parameters at **2 °C/sec ramp rate:** Step 1: 98 °C, 3 min Step 2: 98 °C, 80 sec Step 3: 65 °C, 30 sec Step 4: 72 °C, 60 sec Step 5: Return to Step 2, 14 cycles Step 6: 72 °C, 10 min Step 7: 4 °C, hold
4. If desired, you can run 10 μL of the products of these reactions on a 1.5% agarose gel to confirm the product (195 bp for libraries encoding 11-residue peptides, see **note 5**).
5. Prepare a second primer mix for each sample with the i7/i5 barcode primers, diluting each forward and reverse primer to 5 μM in water. Be sure to use different combinations of barcode primer pairs for each unique sample, as the amplified DNA will eventually be mixed, deep sequenced as a pool, and then de-multiplexed during data analysis based on the unique barcodes.
6. Prepare a PCR reaction for each sample in the following volumes: 5 μL of PCR product from the first reaction 15 μL water 5 μL i7/i5 primer mix 25 μL 2X Q5 master mix
7. Run the PCR according to the following parameters at **2 °C /sec ramp rate:** Step 1: 98 °C, 3 min Step 2: 98 °C, 80 sec Step 3: 65 °C, 30 sec Step 4: 72 °C, 60 sec Step 5: Return to Step 2, 19 cycles Step 6: 72 °C, 10 min Step 7: 4 °C, hold
8. Run the entirety of each PCR reaction on a 1.5% agarose gel, and ***gel purify*** the samples. The expected size is 255 bp for libraries encoding 11-residue peptides. Note that if you see multiple bands, higher than the expected length, you may need to reduce the amount of first-round PCR reaction that you add as a template to the second round PCR reaction.

#### 3.5.2 Quantification of individual DNA samples and preparing the DNA pool for sequencing

1. DNA quantification of the purified samples can be done using the Promega QuantiFluor assay, following the manufacturers protocol. Quantification can also be done using qPCR or another high-fidelity method for quantifying double-stranded DNA. Note that Nanodrop quantification is often not reliable at this stage. Generally, purified DNA samples are in the 5-50 ng/μL range, which corresponds to low to mid nanomolar DNA concentration.
2. Note that the QuantiFluor assay will yield a concentration in units of ng/μL. This can be converted in to nanomolar as follows: [DNA in nM] = ([DNA in ng/μL] / (length in bp x 660)) x 10^6^. Generally, these samples are in the 20 nM range.
3. After quantifying individual samples, prepare a ∼50-100 μL equimolar pool of your samples. Note that if you are conducting screens with libraries of different sizes, adjust your mixture to proportionally overrepresent larger libraries.

### 3.6. Deep sequencing of the pooled samples

Many different deep sequencing platforms can be used to analyze the pooled samples after surface display, phosphorylation, and selection. Thus far, we have primarily used Illumina MiSeq and NextSeq instruments, and conducted paired-end reads using the MiSeq Reagent Kit v3 (150 cycles) or the NextSeq 500 Mid-Output v2 Kit (150 cycles), respectively (see **note 6**). However, we have also successfully analyzed samples prepared as described in **section 3.5** using an Element AVITI sequencer. Here, we briefly describe the protocol for Illumina sequencing of bacterial display samples.

1. In advance of preparing the DNA sample, thaw the Illumina sequencing kit reagent cartridge, following manufacturer’s instructions. Thaw keep the HT1 solution provided with the kit on ice.
2. Dilute your pooled DNA to 4 nM in deionized water. You don’t need to use your whole sample pool, just a few microliters, sufficient to accurately pipette.
3. Thaw the Illumina PhiX control, which should also be at a concentration of 10 nM, and dilute it to 4 nM by mixing 2 μL of PhiX with 3 μL of water.
4. Following the Illumina Denature and Dilute Libraries protocol, mix 5 μL of freshly-prepared 0.2 M NaOH with 5 μL of the 4 nM pooled DNA. Analogously, add 5 μL of freshly-prepared 0.2 M NaOH to the 5 μL of 4 nM PhiX sample. Mix the samples well by vortexing, briefly centrifuge, and incubate them at room temperature for 5 minutes.
5. Next, add 990 μL of thawed HT1 buffer to the pooled sample DNA and to the PhiX tube. The DNA in these tubes should now be 20 pM.
6. Mix the pooled sample DNA and PhiX DNA and further dilute them in HT1 buffer as needed to yield a 600 μL sample that has 8-12 pM total DNA concentration, with 95% pooled sample and 5% PhiX. The exact sample concentration may need to be optimized, to yield an ideal cluster density on a MiSeq chip, but 12 pM is a good starting point for these libraries.
7. Finally, following the Illumina MiSeq or NextSeq user instructions, load your sample onto the cartridge, prepare the instrument, and implement your sequencing run with 75x75 bp paired end reads.

### 3.7 Sequencing data analysis (post-sequencing)

After the samples have been analyzed by deep sequencing, the generated datasets can be analyzed using Python or another desired coding language. This section will assume that FASTQ files containing the raw sequencing reads are available from the sequencing experiment, and that these reads can be processed to obtain translated peptide sequences. These translated sequences can then be used in a variety of calculations to analyze kinase specificity, as described in this section. First, we briefly describe data processing to yield translated sequences, and the methods for generating various counts tables (**section 3.7.1**). The process and Python scripts for generating the counts tables are available on our Github (https://github.com/nshahlab/2022_Li-et-al_peptide-display). While specific procedures for data analysis of generated counts will vary depending on the type of library sorted and desired information, here we describe our standard practices in calculating normalized enrichment scores for head-to-head comparison of different peptides within a given library. First, a general procedure for the calculation of enrichment scores will be described (**section 3.7.2**), followed by specific applications in the context of different libraries (**sections 3.7.3 – 3.7.5**).

#### 3.7.1 Processing the raw sequencing reads

1. First, all of the raw sequencing reads should be transferred to a local folder and decompressed from .fastq.gz to the .fastq file format. There should be a pair of files associated with each barcode pair used in the sequencing run.
2. The matched forward and reverse reads from the paired-end sequencing should be merged. While many programs can be used for this, we often use FLASH (Magoč & Salzberg, 2011).
3. The merged reads should be trimmed. Again, there are various programs to accomplish this, such as Cutadapt (Martin, 2011). Using Cutadapt, all merged sequences containing the following flanks can be identified, and these flanks can be removed, leaving behind only the peptide-coding sequences:

5’ flanking sequence = NNNNNNACCGCAGGTACTTCCGTAGCTGGCCAGTCTGGCCAG

3’ flanking sequence = GGAGGGCAGTCTGGGCAGTCTGGTGACTACAACAAAANNNNNN

1. The trimmed reads can then be translated using a Python script with Biopython modules to generate peptide reads in .fasta files, as described in the aforementioned Github.
2. The resulting translated sequences can then be analyzed using a variety of Python scripts in the Github repository to either determine the counts per unique sequence or the amino acid or stop codon frequency at each position in the peptide.

#### 3.7.2 Calculation of enrichment scores for libraries with discrete peptide sequences

This calculation of enrichment scores is for peptide libraries, such as the 10K phosphosite library, which have a defined set of peptide sequences (**Figure 4**). The protocol assumes that counts for the individual sequences in the library have already been determined for two sequencing files: one from an unsorted sample one from a sample that was phosphorylated and enriched/sorted.

1. Sum the total counts for **all** sequenced peptides in a given sample. These will be “**total counts**.”
2. Divide the counts for each unique peptide sequence by the **total counts**. This will generate a relative “**frequency**” of the specific peptide in a given sample.
3. Repeat steps 1-2 for all samples, including the unsorted control samples which will be the “**unsorted frequency**”.
4. Once all the frequency values have been determined, divide the frequency for each peptide in the sorted samples by the corresponding frequency for that peptide in the matched unsorted sample. This will generate the normalized “**enrichment scores**” for each peptide sequence. These enrichment scores can be used to directly compare peptides to one another, as they correlate strongly with relative phosphorylation rates, as shown previously (Li *et al*, 2023; Shah *et al*, 2016).

**Figure 4.**
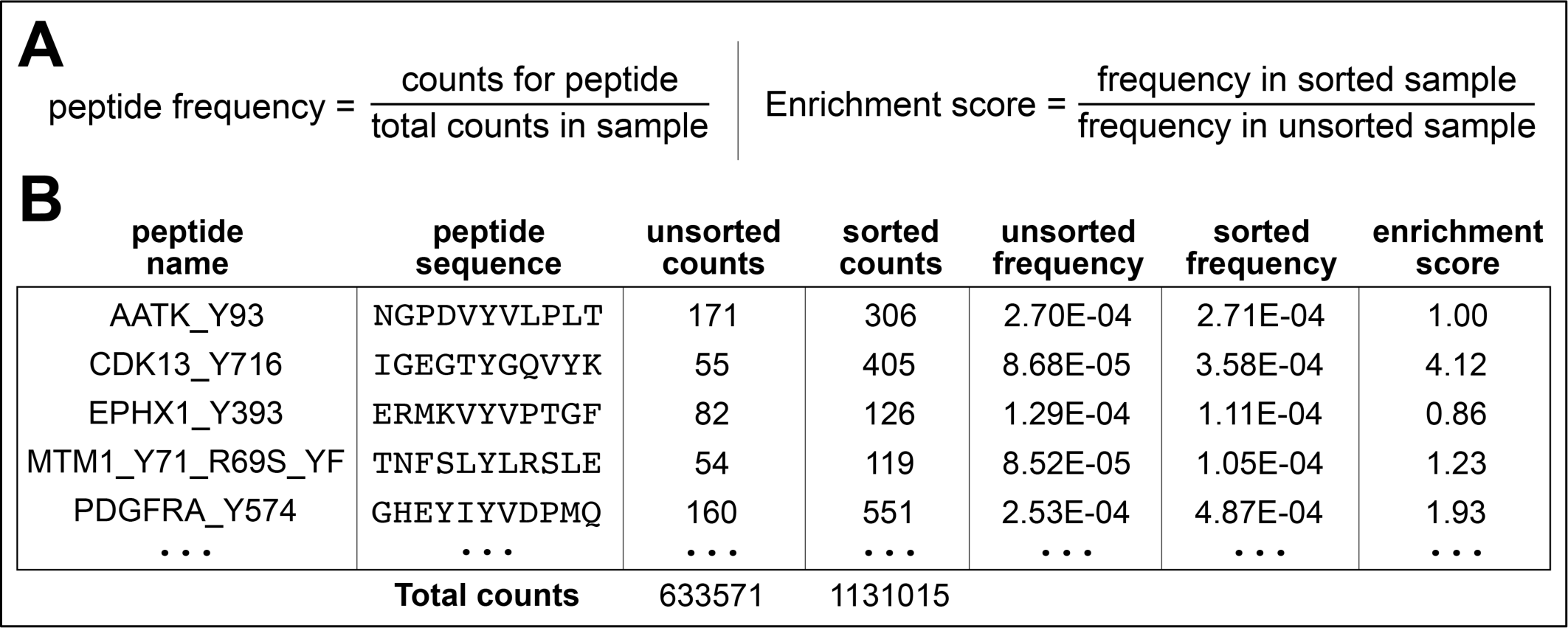
Calculation of enrichment scores for discrete peptide sequences. (**A**) Enrichment scores for each peptide sequence are calculated in two steps: First, the frequency of each peptide is calculated by dividing the counts for a select peptide in a sample by the total counts in that sample. Then, the frequency of a peptide of interest in a sorted sample is normalized to its frequency in a matched sorted sample. (**B**) A segment of a representative data table is shown.

#### 3.7.3 Calculation of enrichment scores for degenerate libraries

For degenerate libraries, which can have >10^6^ different peptides, the frequency of each unique sequence is generally too low to obtain accurate enrichment scores. However, these libraries can be used to readily determine the enrichment of specific amino acids at each position in the sequence. To accomplish this, scripts in the Github can be used to generate a counts matrix that lists the counts for each amino acid at each position along the peptide (**Figure 5A**). These counts matrices from a sorted and unsorted sample can be used to determine the position-specific enrichment of each amino acid (**Figure 5B-D**).

1. Sum the total counts for all amino acids at each position along the peptide sequence length. This will be the **“total sum**” for each residue position relative to the central tyrosine. Repeat this step for all sequenced samples that have degenerate libraries.
2. Divide the counts of each amino acid by the corresponding **total sum** for each position. This will generate a matrix with the relative **“frequencies”** of each amino acid at a given position.
3. Repeat steps 1-2 for all samples, including the unsorted control samples which will be the “unsorted frequencies.”
4. Once all the frequency values have been determined, divide the frequencies for each amino acid at each position in the sorted samples by the corresponding frequencies in the matched unsorted samples. This will generate a matrix of normalized **“enrichment scores”** for each amino acid at a given position in the peptide.
5. These position-specific amino acid enrichments can be log_2_- or log_10_-transformed to readily visualize which sequence features are disfavored (negative), neutral (close to zero), or favored (positive).

**Figure 5.**
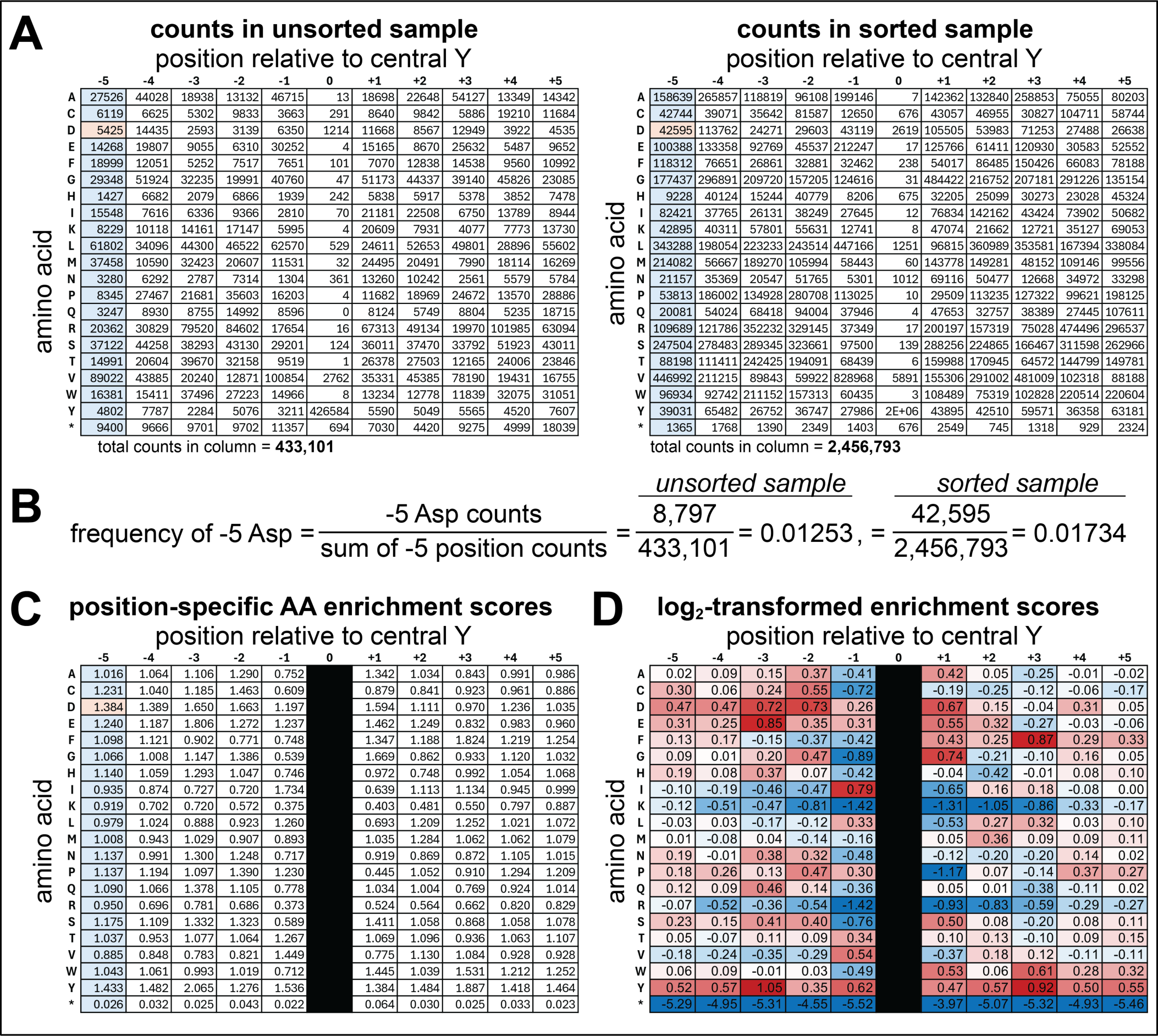
Calculation of position-specific amino acid enrichment scores from a degenerate library. (**A**) Amino acid counts at each position in the X_5_-Y-X_5_ library in an unsorted sample (*left*) and a sample that underwent phosphorylation by c-Src and selection (*right*). (**B**) Example calculation for the frequency of a -5 Asp residue in the unsorted and sorted samples shown in panel (A). (**C**) Position-specific enrichment scores calculated by taking the ratio of frequencies for each amino acid at each position in the sorted and unsorted samples, analogous to how enrichment scores are calculated for individual peptides in Figure 4A. (**D**) Log_2_-transformed position-specific enrichment scores for c-Src specificity against the X_5_-Y-X_5_ library. The normalized scores are colored as a heatmap, with blue indicating a disfavored feature, white being a neutral feature, and red being a favored feature. Note that in panels (C) and (D), the “position 0” column is blacked out, because this library does not encode for a significant number of peptides with residues other than Y at this central position, and thus the enrichment calculations are not accurate or meaningful. In all panels, the row marked by an asterisk indicates introduction of a stop codon at that site in the sequence.

#### 3.7.4 Calculation and normalization of enrichment scores to wild-type for scanning mutagenesis libraries

The calculation of enrichment scores for scanning mutagenesis libraries is similar to how they are calculated for defined sequence libraries, in that the enrichments for specific sequences can be determined. However, for ease of visualization, these data are compiled and represented as a matrix of values representing each point mutant (**Figure 6**). Furthermore, the enrichments for each mutant peptide are often normalized to the reference (wild-type) sequence (**Figure 6**).

1. In the matrix format shown in **Figure 6**, for a scanning mutagenesis library, the reference/wild-type sequence will be repeat-counted in every column, and this should be accounted for in data analysis. Thus, to determine the total counts for a sample, first sum the total counts in the whole matrix. Then, subtract the repeated counts of the reference/wild-type sequence – take the central tyrosine counts and multiply by 10 (or n-1, where n is the total number of residues being scanned in the peptide) to account for the repeated counts. This will be the **“total sum.”**
2. Divide the counts for each amino acid by the **total sum**. This will generate a matrix with the relative **“frequencies”** of each amino acid.
3. Repeat steps 1-2 for all samples, including the unsorted control samples which will be the “unsorted frequencies.”
4. Divide the **frequencies** for each mutant in the sorted samples by the corresponding **unsorted frequencies** in the matched unsorted sample. This will generate a matrix of normalized “**enrichment scores**” for each mutant peptide.
5. Log_2_- or log_10_-transform the enrichment scores, then subtract the log-transformed enrichment value for the reference/wild-type sequence from the scores for all other sequences. This will generate a matrix of log-transformed enrichment scores normalized to the reference/wild-type sequence as a baseline. Here, negative values will indicate mutations that reduce phosphorylation and positive values will indicate mutations that enhance phosphorylation. Mutations at the central Tyr should be significantly negative numbers and serve as an internal negative control.

**Figure 6.**
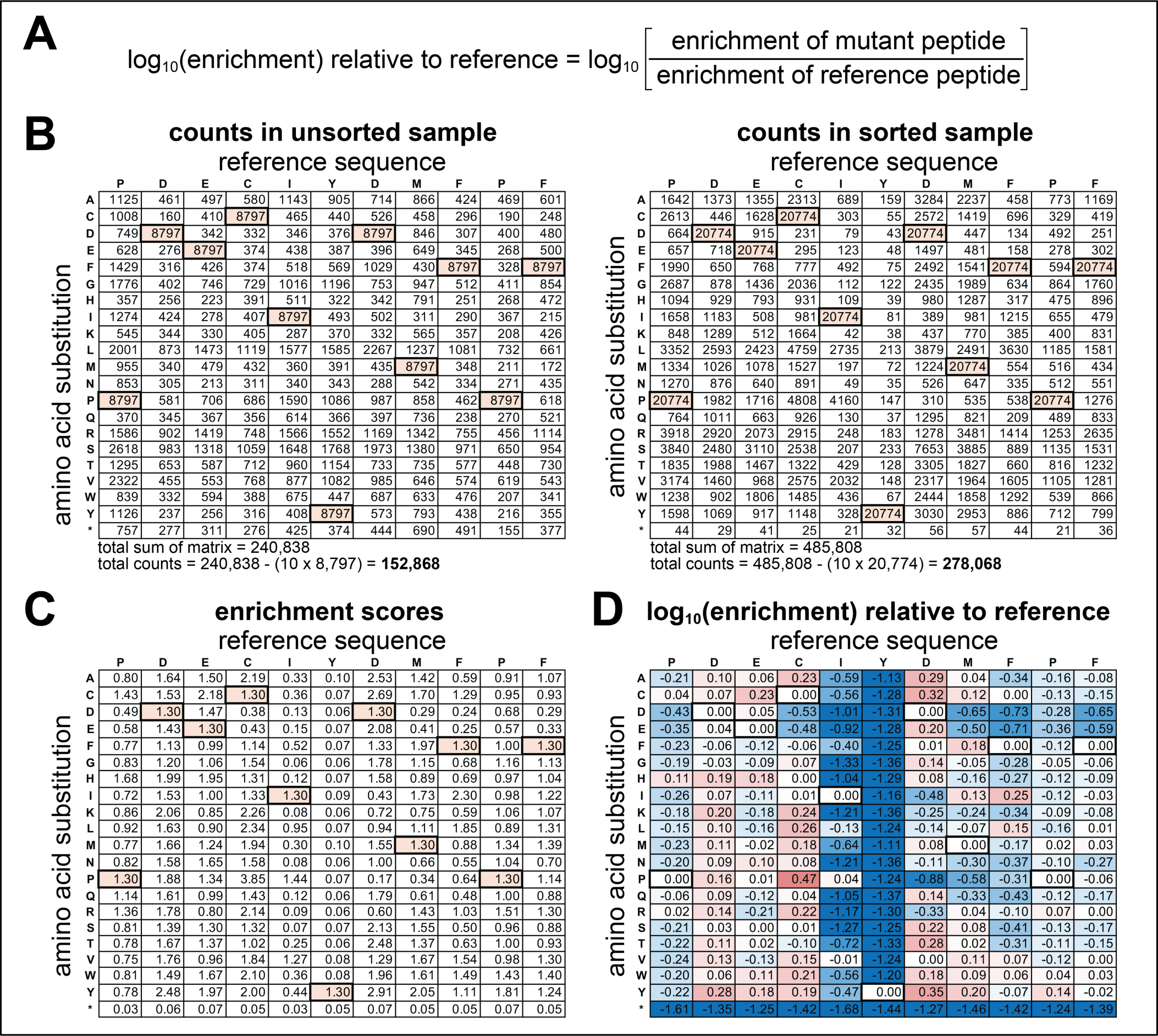
Calculation of normalized enrichment scores in scanning mutagenesis libraries. (**A**) Enrichment scores for individual mutants can be calculated as described in Figure 4A. However, if data are extracted as a mutation matrix, as shown in panel (B), the total counts used to calculate frequencies must account for the repetition of the wild-type sequence in the matrix (bold, colored cells). This is done by taking the total sum of all counts and subtracting the wild-type counts (bold, colored cells) multiplied by 10, or the (sequence length – 1). (**B**) Examples of scanning mutagenesis counts tables for a library based on a high-efficiency Src consensus peptide, phosphorylated by c-Src kinase domain. (**C**) Enrichment scores for all mutants in the scanning mutagenesis library based on the counts in panel (B). (**D**) Log_10_-transformed enrichment scores for all mutants in the library, normalized to the reference sequence, as described in panel (A). The normalized scores are colored as a heatmap, with blue indicating worse than the reference, white being equivalent to the reference, and red being better the reference. In all panels, the row marked by an asterisk indicates introduction of a stop codon at that site in the sequence.

**Figure 7.**
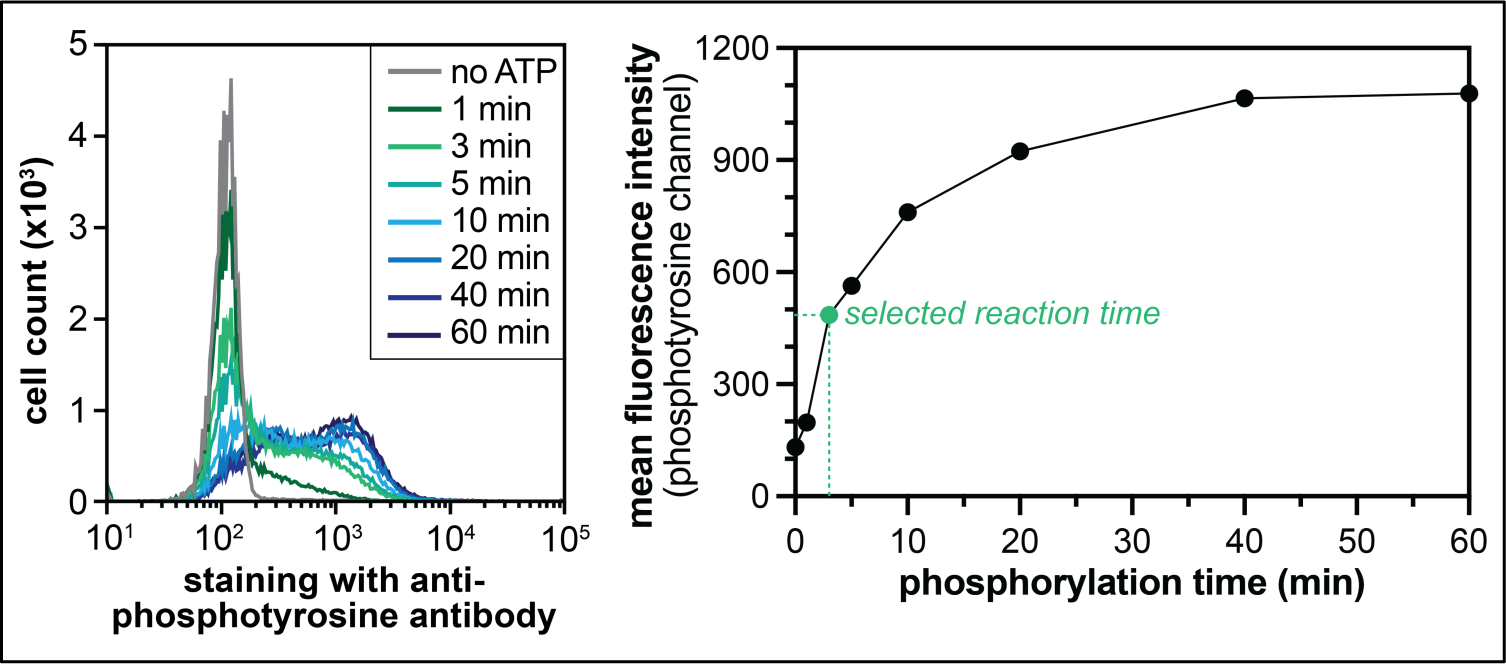
Finding the optimal kinase reaction time. Kinase reaction parameters such as kinase concentration and incubation time can be optimized by measuring phosphorylation over time, staining with a fluorescently-labeled anti-phosphotyrosine antibody, and visualizing phosphorylation levels by flow cytometry (left panel), and quantifying the reaction progress (*right panel*). The data in this figure come from phosphorylating the X_5_-Y-X_5_ library using 500 nM c-Src kinase domain. From this experiment, a time-point of 3 minutes was selected, as the reaction is between 25-50% complete at this time point.

#### 3.7.5 Analysis for amber suppression datasets using a degenerate library

The analysis of datasets involving amber suppression requires additional filtering to exclude sequences that do not contain an Amber stop codon. This is because sequences lacking an internal amber stop codon are expressed with higher efficiency that amber-containing sequences, even when amber suppression is operative. As a result, all amber-containing sequences will look artificially depleted relative to amber-lacking sequences, even if the resulting peptides are actually good substrates (Li *et al*, 2023). Thus, the amino acid counts at each position must be determined only for the subset of sequences that have one amber codon in them. This can be achieved using tailored Python scripts in the Github.

1. As included in the Github, the “**1stop**,” “**full**,” and “**nostop**” Python scripts can be used to generate various counts tables that filter peptides with only one stop codon (and thus only one site of non-canonical amino acid incorporation), total counts (similar to the previously described analyses), and filter peptides with no stop codons. The amber suppression datasets are analyzed using the “**1stop**” scripts to filter for peptides that have exclusively one amber codon, and thus one non-canonical amino acid.
2. To analyze data from kinase specificity screens with amber suppression, repeat the steps described in **section 3.7.3** to generate the position-specific enrichment scores all canonical amino acids and for the non-canonical amino acid (represented as enrichment of the stop codon).

## 4. Notes

1. The resuspended cells in cold D-PBS can be stored for up to one week at 4 °C, from our experience.
2. Kinase reaction parameters (incubation time, kinase concentration) can be optimized by monitoring the extent of library phosphorylation using flow cytometry. Cells can be phosphorylated for varying amounts of time, with different kinase concentrations, and quenched, as described in **section 3.4**. Then, instead of labeling with a biotinylated pan-phosphotyrosine antibody, a fluorescently-labeled antibody can be used. The cells are then analyzed by flow cytometry to identify conditions where the extent of phosphorylation is between 25-50%, which is a good range for library selections (**Figure 7**).
3. Working volume for selection can be halved, to save on beads. The protocol described specifies 75 μL of beads/sample, but this can be halved to 37.5 μL beads/sample if using 50 μL of cells instead of 100 μL.
4. Make sure the beads are all settled to the bottom of the microcentrifuge tube before boiling for lysis (none stuck on the sides).
5. A 1.5% agarose gel may be run on 10 μL of each sample after the first PCR (TruSeq primer attachment step), if desired. We have use gel imaging software to quantify the bands, if visible, to estimate volumes required for the second PCR. It is noteworthy that the bands will not always be clear after this first PCR.
6. The scale of sequencing and the number of samples pooled should be determined based on the desired depth. In general, for defined libraries, we want at least 200, but up to 500, reads for each sequence, in an unsorted sample. Thus, for the 10K phosphosite library, one sample will require at least 2 million reads. A MiSeq v3 kit can produce 20-30 million reads, which means that 10-15 samples can be multiplexed. For the degenerate X_5_-Y-X_5_ library, it is not practical to sequence with that level of depth, given the library complexity, so we typically aim for 1-2 million reads per sample using this library.

## 5. Acknowledgements

We thank members of the Shah lab for their valuable contributions throughout the development of this method. This research was funded by NIH/NIGMS grant R35GM138014 and NSF CAREER Award 2441001 to NHS.

